# An unexplored diversity for adaptation of germination to high temperatures in *Brassica* species

**DOI:** 10.1101/2024.10.22.619588

**Authors:** M. Tiret, M.-H. Wagner, L. Gay, E. Chenel, A. Dupont, C. Falentin, L. Maillet, F. Gavory, K. Labadie, S. Ducournau, A.-M. Chèvre

## Abstract

Elevated temperatures inhibit the germination of a concerning number of crop species. One strategy to mitigate the impact of warming temperatures is to identify and introgress adaptive genes into elite germplasm. Diversity must be sought in wild populations, coupled with an understanding of the complex pattern of adaptation across a broad range of landscapes. By investigating the landraces, wild, and feral populations of Algeria, Italy, France, Slovenia, Spain, and Tunisia, we assessed the response of germination to temperature increase in an unexplored diversity of 117 accessions of *B. rapa* and 66 of *B. oleracea*. Our results show that both species exhibit heat tolerance to the temperature range tested, especially *B. rapa* with an increase in speed and uniformity of germination time, as well as an increase in germination rate as temperature increased. As for *B. oleracea* accessions, the ability to germinate under heat conditions depended on the geographical origin; in particular, southern populations showed a higher germination rate than northern populations, possibly in relation to their warmer climates of origin. These findings highlight the complex interplay between domestication, feralization, and current agronomic practices in shaping germination characteristics in *Brassica* species.

## Introduction

Environmental factors, including temperature, precipitation patterns, increased atmospheric CO2 concentrations, and extreme weather events, will undergo significant deviations from their current states as a direct consequence of climate change (IPCC, 2013; IPCC, 2021). Agriculture is among the most endangered sector (Wheeler & Von Braun, 2013; Arunanondchai et al., 2018; Raza et al., 2019), across many major regions (Rosenzweig et al., 2014; Thornton et al., 2014), particularly due to warming temperatures often leading to reduced yields, altered crop quality, and increased susceptibility to environmental variations (Lobell & Field, 2007; DaMatta et al., 2010; Zhao et al., 2017; Chaudhry et al., 2022). On many major crops, the detrimental effects of warming temperatures are evident in the disruption of developmental stages and phenological cycles (Angadi et al., 2000; Barnabás et al., 2008; Chmielewski et al., 2004; Hatfield & Prueger, 2015).

Seed vigor is particularly vulnerable to the environmental changes that are induced by climate change (Hampton et al., 2016; Castillo-Lorenzo et al., 2019). Seed vigor encompasses seed longevity, germination speed, seedling growth, and early stress tolerance (Pollock & Roos, 1972; Reed et al., 2022). It is a key determinant of a plant’s early growth and competitive ability, and is essential for ensuring successful establishment (TeKrony & Egli, 1991; Finch-Savage & Bassel, 2016; Ebone et al., 2020). The processes involved are influenced by a variety of environmental factors, either experienced during seed development (i.e., the maternal effect; Wolf & Wade, 2009; Penfield & MacGregor, 2017), storage (Taylor, 2020), or on the seedbed (Lamichhane et al., 2018). Elevated temperatures are known to impair seed fertility, delay germination, and lower seed vigor in various crop species (Gareca et al., 2012; Cochrane et al., 2015; Tribouillois et al., 2016; Reed et al., 2022).

The susceptibility of seed vigor to environmental factors has often been ascribed to evolutionary responses to specific geographical conditions (Clauss & Venable, 2000; Galloway, 2005; Franks et al., 2014; Dürr et al., 2015; Becklin et al., 2016). Adaptation to the local environment is a widespread phenomenon, as plants that evolved in warmer climates often have the ability to thrive in high-temperature conditions (Cochrane et al., 2015; El-Keblawy et al., 2018). However, the recent ongoing climate change may require a shift in phenotypes to remain locally adapted, but whether rapid genetic change is sufficient to keep pace with climate change remains a topic of debate (Jump & Peñuelas, 2005; Jump et al., 2008). The relationship between climate of origin and heat tolerance is complex and it is not always straightforward to identify a pattern of local adaptation due to phenotypic plasticity and the interplay of various environmental and genetic factors (Jump & Peñuelas, 2005; Nicotra et al., 2010). These studies underscore the intricate nature of a ‘climate-smart agriculture’ strategy to mitigate the impact of climate change (Pareek et al., 2020; Sloat et al., 2020).

The *Brassica* genus is notable for its extensive morphological and genetic diversity, which has made it a global economic resource, particularly in the production of edible root crops, vegetables, and oilseeds (Warwick & Black, 1991; Rakow, 2004; Cheng et al., 2016). Despite this diversity, cultivated lines exhibit limited genetic variation, particularly those of oilseed rape (*Brassica napus* L.), which constrains their adaptive potential (Allender & King, 2010). Establishment failure has been identified as a recurring problem for oilseed rape, and despite extensive research, a solution remains elusive (Nelson et al., 2022). As in many plant genera, the utilization of plant genetic resources in the *Brassica* genus has been considered a strategy to mitigate climate change (Salgotra & Chauhan, 2023). Wild populations and crop wild relatives represent an untapped gene pool, adapted to a wide range of habitats and potentially more resilient to climate change than cultivated crops due to their higher genetic diversity (Castillo-Lorenzo et al., 2019; Renzi et al., 2022). They offer valuable traits for improving stress tolerance and adaptability to changing environmental conditions, making them prime candidates for breeding programs (Tanksley & McCouch, 1997; Zamir, 2001; Nevo et al., 2012; McCouch et al., 2013). In this regard, crop wild relatives of the *Brassica* genus have been screened recently for heat and water stress resistance (Castillo-Lorenzo et al., 2019). Nevertheless, a large-scale sampling strategy across a broad range of climatic conditions remains to be conducted. The objective of this study was to assess, with high-throughput imaging phenotyping, the germination abilities under heat stress conditions of an unexplored diversity panel of wild populations and landraces comprising 117 accessions of *Brassica rapa* L. and 66 accessions of *Brassica oleracea* L., the two progenitors of *B. napus*, and provide insights for future targets of breeding aiming to mitigate the effect of climate change.

## Materials & Methods

### Plant material

The material, which is a subsample of the plant collection presented in Falentin et al. (2024), comprises 117 accessions of *B. rapa* and 66 of *B. oleracea*. Among them, 55 accessions of *B. rapa* and 50 of *B. oleracea* are landraces, and 62 accessions of *B. rapa* and 16 of *B. oleracea* are populations sampled from non-cultivated areas. For simplicity, we will refer to these populations as ‘wild’ in the following, knowing that it may reflect different origins (i.e., true wild or feral populations; see Mittel et al., 2020; McAlvay et al., 2021; Saban et al., 2023). Seeds were collected in Algeria, France, Italy, Slovenia, Spain, and Tunisia. For wild populations, seeds were collected from a maximum of 30 mother plants per sampling site; for landraces, they were collected from local farmers. To circumvent any potential maternal effect, the collection was then multiplied over two consecutive years (2020/2021 and 2021/2022) at a single location (Le Rheu, France, 48°06’22”N 1°47’25”W). Seedlings were grown for seed production under cages covered with pollen-proof netting: two cages per population of *B. rapa* with five plants each, and three cages per population of *B. oleracea* with two plants each. The production of the multiplication is what we will refer to as the “accession”, which is a heterogeneous pool representing the descent of 6 to 10 mother plants (Table 1). Hereafter, we refer to the accessions’ affiliation with wild populations or cultivated landraces as their ‘type’. Twenty-eight accessions of *B. rapa* and 50 of *B. oleracea* were harvested in the summer of 2021, and 107 accessions of *B. rapa* and 16 of *B. oleracea* were harvested in the summer of 2022 (18 accessions of *B. rapa* overlapped between the two seasons). Seed lots were stored at 4°C and 10 % relative humidity in the dark until phenotyping. Seed lots were then split for different analyses and measurements (germination in controlled conditions and emergence in field conditions).

**Table 1.**
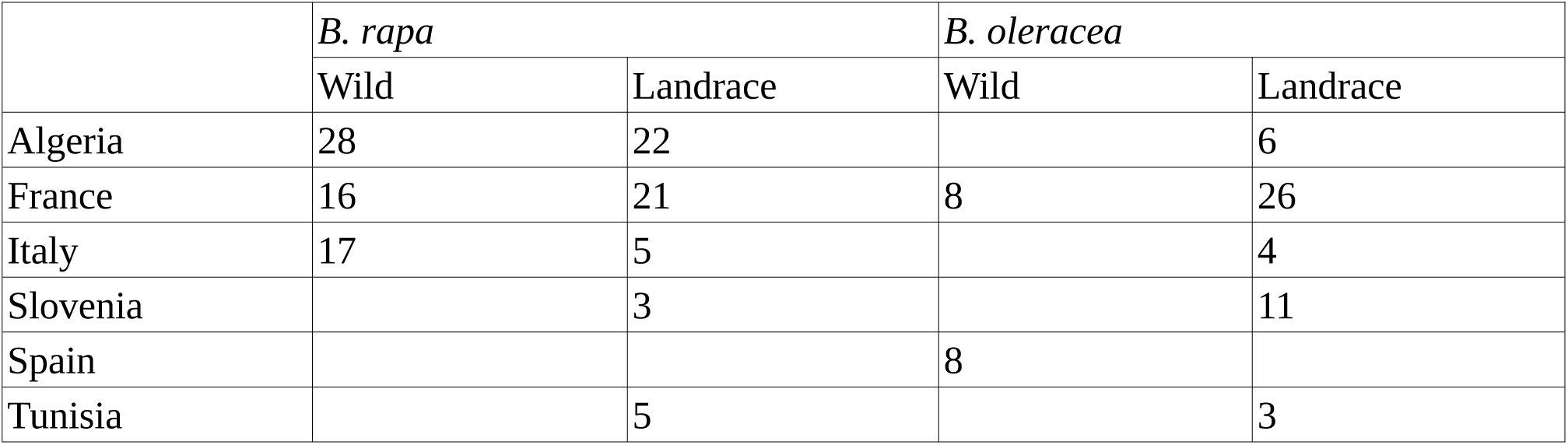
The accessions by types (wild or landrace) and countries (Algeria, France, Italy, Slovenia, Spain, and Tunisia), out of the 117 accessions of *B. rapa* and 66 accessions of *B. oleracea*.

### Phenotyping – seed germination

We phenotyped the accessions for germination time (GT) and rate (GR) under two treatments in controlled conditions: at 20°C following the ISTA reference temperature for both species (ISTA, 2024), and at 35°C for *B. rapa* and 30°C for *B. oleracea* to measure heat tolerance. The maximum temperatures were determined based on tests on commercial germplasm, on what was perceived as a stress (germination rate drops).

Germination was monitored using the high-throughput Plant Imaging PHENOTIC platform at the National Seed Testing Station at Angers, France (GEVES, Angers, France). Measurements were performed on 16 trials of eight days on Jacobsen germinators (paper support), from 15 September 2021 to 6 June 2023. Seeds were imbibed in the dark for 168 h at a water potential measured to be −0.4 MPa (due to the water retention capacity of the thick germination paper and the capillarity equilibrium of the incubator), which is out of the range of drought stress. During the experiment, the germinators maintained a constant temperature of 20°C or 30/35 ± 5°C. The germination process (radicle emergence) was automatically assessed via the image analysis method described in Demilly et al. (2014), with images captured every two to three hours throughout the experiment. When necessary, the data collection was terminated after seven days to minimize the occurrence of false positive. At the end of the experiment, if germination occurred within the first 168 hours, the time to the start of germination (in hours), namely GT, was recorded; otherwise, it was declared as non-germinated.

In each trial, a randomized incomplete block design was implemented, with blocks of 25 seeds of a single accession (one batch; see details in Wagner et al., 2011). All accessions were replicated at least 100 times (four batches) per treatment (low or high temperature), with the exception of four accessions of *B. rapa* replicated 95 times at 35°C, all due to material availability. Additionally, to investigate the effect of the seed multiplication year, supplementary seeds of *B. rapa* were tested based on their availability, leading to an additional 25 seeds for 22 accessions, 100 seeds for 34 accessions and 125 seeds for 10 accessions at 20°C, and an additional 100 seeds for five accessions at 35°C. For *B. rapa*, only 79 accessions (out of 117) were tested at 35°C given that the germination rate was already low at 20°C: temperature increase would not have broken dormancy, which is the major reason of low germination at 20°C for Algerian and Italian wild accessions. For *B. oleracea*, all 66 accessions were tested at both temperatures with 100 seeds each, with the exception of four accessions that were tested at 30°C with 125 seeds. In total, 25,280 seeds of *B. rapa* (16,900 at 20°C and 8,380 at 35°C) and 13,300 of *B. oleracea* (6,600 at 20°C and 6,700 at 30°C) were phenotyped.

### Phenotyping – seed emergence

To confirm that the germination ranking is maintained under field conditions, we conducted an experiment in a common garden setting. The emergence rate, defined as the ratio of cotyledon-stage seedlings to the total number of seeds sown, was measured in the field in 2022 on the same lots of seeds that were used in the laboratory. For the *B. rapa* accessions, the seeds were sown in the field in a random incomplete block design on 10 September 2022 at Le Rheu (France, 48°06’33”N 1°47’11”W; with an average daily temperature of 19.3°C during the following week), in three repetitions of 120 blocks (including a control replicated three times in each repetition, the “Purple Top Milan”), with 150 seeds per plot of 1 m x 1.5 m. For the *B. oleracea* accessions, the seeds were sown in small pots within a greenhouse between 19 September 2022 and 22 September 2022, at Ploudaniel, France (48°30’06”N 4°19’32”W; with an average daily temperature of 18°C during the following week), with 140 seeds per accession. The emergence rate was measured on a block on 6 October 2022 for *B. rapa* and on 3 October 2022 for *B. oleracea*.

### Sequence data and bioinformatics

For each accession, a pool of 30 seedlings (for landrace accessions, 30 random seeds; for wild accessions, at least one seed per mother plant) was used to generate DNA bulks with a standardized sampling per plant (see details in Maillet et al., 2023). Libraries were constructed from the DNA pools (250 ng) using the NEBNext DNA Sample Prep modules (New England Biolabs) with ‘on beads’ protocol. DNA libraries were sequenced using Illumina NovaSeq 6000 technology (short reads, paired-end, 150 bp; Illumina Inc., San Diego, CA) with a target coverage of 50X. The sequences of the Illumina adapters and primers used during library construction were removed from the whole reads. Low quality nucleotides with quality value < 20 were removed from both ends. The longest sequence without adapters and low quality bases was kept. Sequences between the second unknown nucleotide (N) and the end of the read were also trimmed. Reads shorter than 30 nucleotides after trimming were discarded. These trimming steps were achieved using fastx_clean (http://www.genoscope.cns.fr/fa), an internal software based on the FASTX library (http://hannonlab.cshl.edu/fast). The reads and their mates that mapped onto run quality control sequences (Enterobacteria phage PhiX174 genome) were removed using SOAP aligner (Li et al., 2008). The cleaned reads were mapped on the *B. rapa* ‘C1.3’ var. *rapifera* genome for *B. rapa* (which corresponds to the A subgenome of the *B. napus* ‘RCC-S0’; Maillet et al., 2023) and *B. oleracea* ‘HDEM’ ssp. *capitata* genome for *B. oleracea* (Belser et al., 2018), using the default parameters of the ‘mem’ program from the bwa v0.7.17 software (Li & Durbin 2009; Li, 2013). Read alignments with a mapping quality Phred score below 20 or PCR duplicates were then removed using the view (option ‘-q 20’) and ‘markdup’ programs from the SAMtools v1.14 (Danecek et al., 2021) software suite. Then, we completed the mapping by a step of realignment around indels for *B. oleracea*, as implemented in GATK (McKenna et al., 2010). Subsequently, variant calling was conducted using BCFtools v1.11 (Danecek et al., 2021) with the options ‘-q 20 - Q 20’, which set the minimum mapping quality for an alignment and the minimum base quality for a base to 20.

The resulting VCF file (containing 12,771,634 SNPs in *B. rapa* and 10,778,346 SNPs in *B. oleracea*) was filtered using BCFtools to retain only biallelic SNPs, and remove SNPs situated less than 5 bp away from indels. Only accessions with an average depth higher than 20 were retained (removing five accessions of *B. rapa* and six of *B. oleracea*). Then, SNPs were retained if they exhibited an average depth between its 5th and 95th percentiles, a mapping quality exceeding 40, and mapped on chromosome scaffolds. SNPs were also removed when both allelic variants had a depth below 10 to avoid false positive in downstream quantitative genetics analyses; this filter also removed all SNPs with missing values. The VCF file was then pruned for linkage disequilibrium, with the prune program from the BCFtools v1.11 software suite, with the option ‘-l 0.2 -w 1000’. The variant filtering resulted in 106,064 SNPs for the 112 accessions of *B. rapa*, and 80,054 SNPs for the 60 accessions of *B. oleracea*.

### Population genetic analyses

The genomic relationship matrix (G) was calculated using GCTA v1.94.1 (Yang et al., 2011) with the default parameters, and subsequently transformed into a dissimilarity matrix (1-G*), where the correlation matrix (G*) was derived from G using the ‘cov2cor’ function of the R package ‘stats’. The dissimilarity matrix was analyzed using a multidimensional scaling (MDS) decomposition with the ‘cmdscale_lanczos’ function of the R package ‘refund’ (Goldsmith et al., 2024). The subpopulation membership was calculated using the ADMIXTURE program v1.22 (Alexander et al., 2009), with the default parameters and a cross-validation (CV) error rate used to determine the likelihood of the K parameter.

### Statistical analyses

All analyses were conducted with R v4.4.1 (R Core Team, 2024). The genomic Best Linear Unbiased Predictor (gBLUP) of germination time (GT) for the accessions was calculated using a mixed linear model (one for each species) according to the following model:

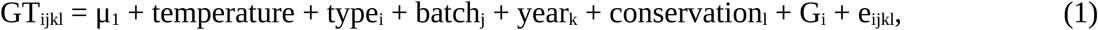

where ‘GT_ijkl_’ is the germination time of the i-th accession, in the j-th batch, of the k-th seed lot, of the l-th seed. The factor ‘μ_1_’ is the intercept, ‘temperature’ the fixed effect of the heat treatment, ‘type’ the fixed effect of the type of the accession (wild or landrace), ‘batch’ the fixed effect of the batch in the germination experiment, ‘year’ the fixed effect capturing the environmental conditions of the year of multiplication (2020/2021 or 2021/2022), and ‘conservation’ the fixed effect of the number of days the seeds had been conserved before phenotyping (controlling for the detrimental effect of seed aging). ‘G’ is the random polygenic effect following a normal distribution of mean 0 and variance matrix σ_G_^2^G, where σ_G_^2^ is the additive genetic variance, and ‘e’ is the residual, also following a normal distribution of mean 0 and variance matrix σ_e_^2^I, where σ_e_^2^ is the residual variance. In accordance with the recommendations set forth by Crawley (2007), we also fitted the model to subdivided datasets: separating wild populations and landraces (two models denoted 1b), and separating wild and landraces for each temperature (four models denoted 1c). The model was fitted using the function ‘remlf90’ of the ‘breedR’ package (Muñoz & Sanchez, 2024), with the EM method, and one step of the AI method to estimate the standard error.

For the proportion of seeds that germinated, hereafter called the germination rate (GR), it was first corrected for residual factors, a process analogous to the approximated method implemented in GRAMMAR as a pre-treatment for binary variables for computing gBLUP (Aulchenko et al., 2007):

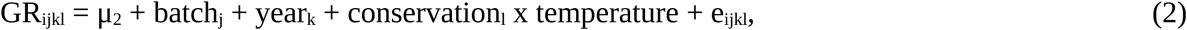

where ‘GR_ijkl_’ is the indicator (0 or 1) of whether the seed germinated before 168 h, ‘μ_2_’ is the intercept, ‘conservation x temperature’ the fixed interaction effect between conservation length and temperature, with the indices and the other factors defined as above. Model (2) was fitted to a binomial distribution using the function ‘glm’ of the ‘stats’ package. The residues of the fitted model (2) (in the latent scale), thereafter called ‘GRA’, were then fitted with the following mixed linear model:

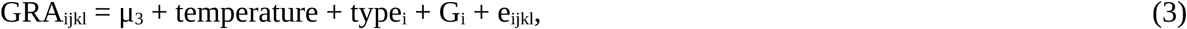

where ‘GRA_ijkl_’ is the adjusted germination rate, ‘μ_3_’ is the intercept, with indices and factors defined as above. Once more, the model was fitted to subdivided datasets: separating wild and landraces (two models denoted 3b), and separating wild and landraces for each temperature (four models denoted 3c). All estimations of GR were conducted on the latent scale. The decision to employ two successive models ensures reliable convergence of the estimation process, which is not currently guaranteed by any available software implementing a (Bayesian) all-in-one GLM approach for computing the gBLUP, including tools like the ‘BGLR’ package (Pérez & de Los Campos, 2022) that offer robust methods but require separate model specification. The effect of temperature was tested using a Z-test (which provides a reliable approximation of the Student’s t-test since the sample size is large), and the effect of the type was tested with a Tukey’s HSD test. Heritability, defined as σ_G_^2^/(σ_G_^2^ + σ_e_^2^), was estimated with models (1c) and (3c). To refine the trend by the contrasting response of different countries of origin, we also considered the following model:

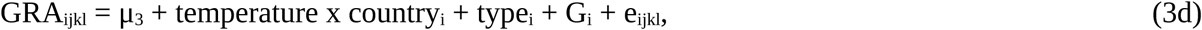

where “temperature x country” is the per-country slope of temperature, with indices and factors defined as below. For each species, the gBLUP from the models (1c) and (3c) of the different temperatures were compared throughout the following model:

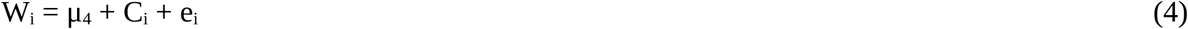

where ‘W_i_’ is the gBLUP at warm temperature, C_i_ the gBLUP at cold temperature, ‘μ_4_’ is the intercept, with indices and factors defined as above. The GPS coordinate of the sampling site (available in Tables S2 and S3 of Falentin et al., 2024) was used to extract the bioclimatic variables from WorldClim v2.1 (climate data for 1970-2000; Fick & Hijmans, 2017) using ‘geodata’ (Hijmans et al., 2024) and ‘terra’ (Hijmans, 2024) R packages, with an accuracy of ten minutes of degree. The Pearson’s correlation coefficient between the latitude and the ‘bio1’ bioclimatic variable (annual mean temperature) was found to be of high amplitude (−0.89), so latitude was used as a proxy for bioclimatic variables in the subsequent analyses. For each species, we studied the link between the latitudes of the sampling sites and heat tolerance H_i_, defined as W_i_-C_i_ for the i-th accession, with the following model:

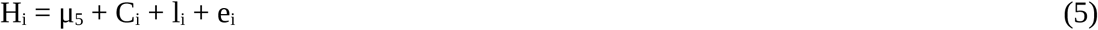

where ‘H_i_’ is the heat tolerance of the i-th accession, ‘l_i_’ is the latitude at the sampling site, ‘μ_5_’ is the intercept, with indices and factors defined as above. ‘C_i_’ was added as an explanatory factor to control for the baseline effect. Finally, we tested the relationship between the gBLUP of GR from model (3c) and the emergence rate by fitting the following model for each species:

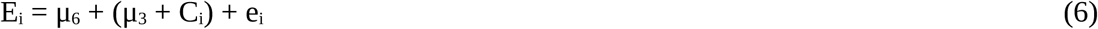

where ‘E_i_’ is the emergence rate, ‘μ_3_’ is the intercept of the model (3c) for cold temperatures of the corresponding species, ‘μ_6_’ is the intercept, with indices and factors defined as above.

## Results

### 1. The type of the accession was a reliable predictor of its germination time and rate

The genetic divergence among the accessions was mainly explained by the wild-landrace axis (Fig. 1A and 1D). In *B. rapa*, Algerian and Italian wild accessions exhibited stronger divergence from the landraces and wild French accessions. In both MDS and admixture analyses (best K = 3 from CV), French accessions clustered with landraces. The differentiation between wild and landraces was carried by the first axis of the MDS, which explained 77.88% of the variance. In *B. oleracea*, however, this distinction was not as clear as for *B. rapa*: MDS and admixture analyses (best K = 4 from CV) partially differentiated wild and landraces (48.07% in the first axis). The wild accessions were closest to *B. oleracea* ssp. *acephala* and *B. oleracea* ssp. *medullosa* (Fig. S1).

**Figure 1.**
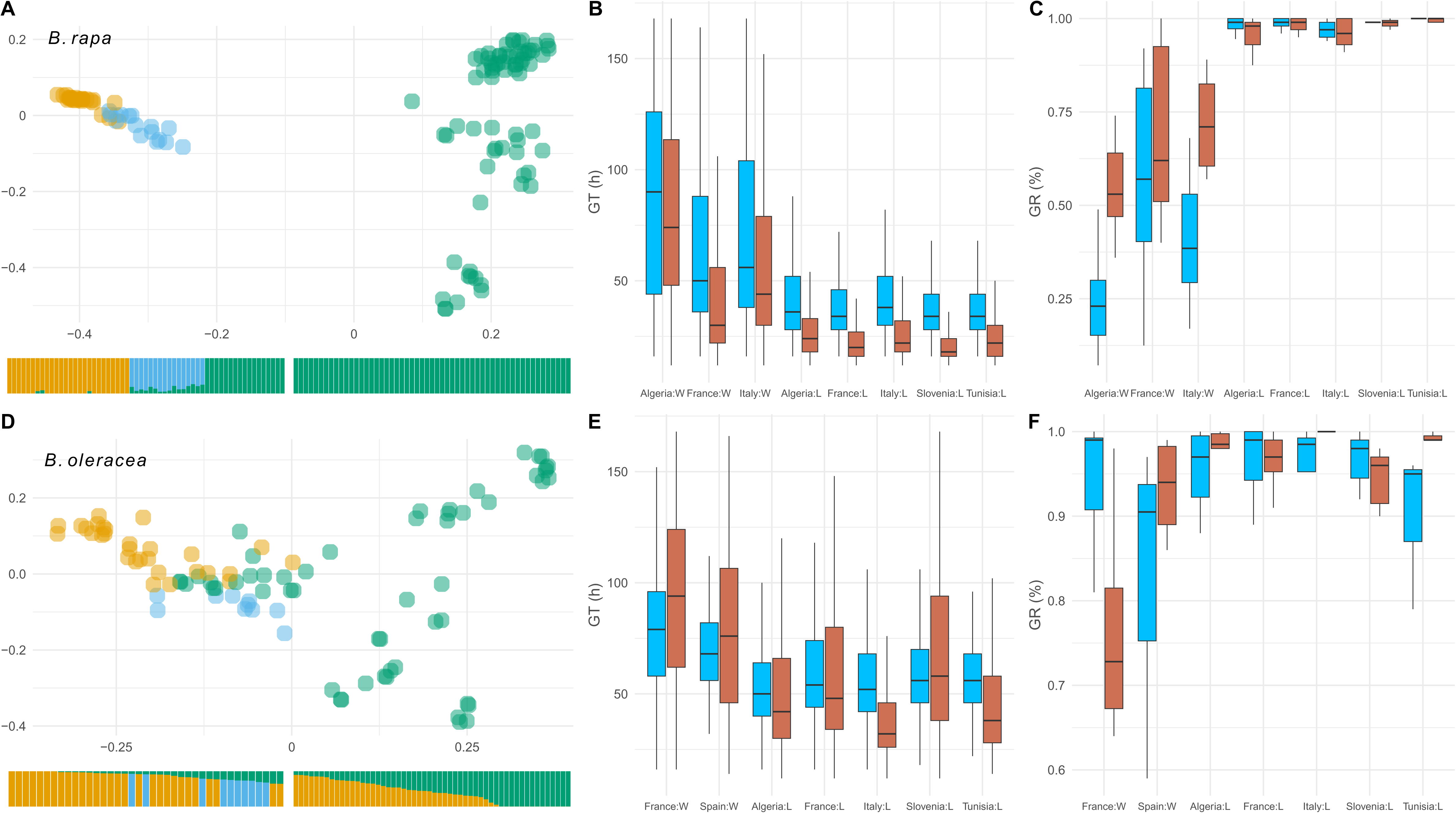
Genetic clusters and germination of Brassica wild and landrace accessions. **(A)** MDS of the genotypes of *B. rapa*: the first axis explains 77.58 %, and the second 12.11 %. Admixture plot in the bottom with the corresponding color (Algerian wild accessions in orange, Italian wild accessions in blue, landraces in green; green wild accessions on the left panel is French landraces). **(B)** Germination time (in hours) of *B. rapa*: 20°C in blue, 35°C in red. **(C)** Germination rate (in percent) of *B. rapa*: 20°C in blue, 35°C in red. **(D)** MDS of the genotypes of *B. oleracea*: the first axis explains 48.07 %, 23.23 %. Admixture plot in the bottom with the corresponding colors (French wild accessions in orange, Spanish wild accessions in blue, landraces in green; Spanish accessions have been colored post-analysis). **(E)** Germination time (in hours) of *B. oleracea*: 20°C in blue, 30°C in red. **(F)** Germination rate (in percent) of *B. oleracea*: 20°C in blue, 30°C in red.

Similar to the genetic divergence between wild and landrace accessions, there was a clear distinction regarding the germination pattern (Table 2). The type of accession (wild or landrace) was the most discriminating factor: germination time (GT) was significantly longer for wild accessions than for landraces in the case of *B. rapa* (Tukey’s test, q = 12.80, p < 0.001; model 1) and of *B. oleracea* (q = 7.98, p < 0.001; model 1), with a slowing effect of 25.92h and 21.82h, respectively. Similarly, germination rate (GR) was significantly lower for wild accessions in both species (q = −15.05 and −5.73, p < 0.001 and < 0.001 for *B. rapa* and *B. oleracea* respectively; model 3). *B. oleracea* accessions were slower than *B. rapa* to germinate, regardless of the type or condition (Student’s t-test, t > 19.56, p < 0.001), except for wild accessions at 20°C (t = 0.61, p = 0.540). The heritability of GT and GR was intermediate (Table S1), with an average of 21.25% for GT (ranging from 13 to 34%) and of 14.00% for GR (ranging from 8 to 22%). As expected, the additive variances are systematically larger in wild accessions than in landraces (Table S2), in particular for GR. The genome-wide association studies (GWAS) on germination time and rate across temperatures did not identify any major genes (Fig. S2-S9; see section A of Supplementary Materials).

**Table 2.** Average germination rate (before 7 days) and germination time (in hours), and standard deviations, of *B. rapa* and *B. oleracea* in controlled conditions (20°C and 35°C for *B. rapa*, 20°C and 30°C for *B. oleracea*).

|  | T° | *B. rapa* |  | *B. oleracea* |  |
| --- | --- | --- | --- | --- | --- |
|  |  | Wild | Landrace | Wild | Landrace |
| GR | 20°C | 0.39 ± 0.49 | 0.97 ± 0.16 | 0.88 ± 0.32 | 0.96 ± 0.20 |
|  | 30/35°C | 0.67 ± 0.47 | 0.97 ± 0.18 | 0.84 ± 0.37 | 0.96 ± 0.20 |
| GT | 20°C | 74.0 ± 41.9 | 42.6 ± 23.9 | 74.5 ± 23.6 | 59.7 ± 24.0 |
|  | 30/35°C | 56.6 ± 40.4 | 27.0 ± 18.7 | 85.9 ± 40.0 | 57.8 ± 33.7 |

### 2. Elevated temperature altered germination of *B. oleracea* accessions, but enhanced that of *B. rapa* accessions

In *B. rapa*, GR was not impacted by the year of production (Z-test, z = −1.74, p = 0.08), but was by the conservation time, where GR was significantly lower for those conserved for a long period of time (z = −21.63, p < 0.001; model 2). In *B. oleracea*, GR was affected by both the year of production (z = −4.75, p < 0.001; model 2; with higher GR for 2020/2021) and conservation time (z = −10.55, p < 0.001; model 2; lower GR when conserved for a long period of time). Models (1) and (3) account for these effects.

Accessions of *B. rapa* germinated faster when temperature increased, for both wild and landrace accessions (z < −7.36, p < 0.001; model 1b; Fig. 1B), with a reduction of 0.64 h/°C and 1.60 h/°C respectively. Temperature slightly but significantly increased GR for both landrace accessions (z = 29.13, p < 0.001; model 1c) and wild accessions (z = 16.93, p < 0.001; model 1c; Fig. 1C), with an effect size three time larger for wild accessions. The trend is consistent across countries for landraces (z > 6.76, p < 0.001; model 3d), yet less pronounced for wild accessions: the effect sizes of wild Algerian and Italian accessions were four times larger (z > 13.10, p < 0.001; model 3d) than French wild accessions (z = 5.62, p < 0.001; model 3d). In contrast, in *B. oleracea* (Fig. 1E), wild accessions germinated significantly slower when temperature increased (z = 9.56, p < 0.001; model 3b), adding 1.31 h/°C, but not for landrace accessions (z = −0.83, p = 0.405). In terms of GR, a slight but significant impact was observed in landrace accessions only: an increase for landraces (z = 5.60, p < 0.001; model 3b), mainly driven by Algerian, Italian and Tunisian accessions (z > 4.60, p < 0.001; model 3d) with an effect size two times larger than the slight increase in French accessions (z = 3.70, p < 0.001; model 3d) or even no effect in Slovenian accessions (z = 1.19, p = 0.234; model 3d). No discernible trend emerges for wild accessions (z = −1.33, p = 0.185; model 3b), due to heterogeneous trends wherein: French wild accessions showed a decrease in GR when temperature increases (z = −6.96, p < 0.001; model 3d), while Spanish accessions showed an increase in GR (z = 5.07, p < 0.001; model 3d).

### 3. Ranking was maintained across temperatures for germination time, but not for germination rate

It would be interesting to know whether the significantly different abilities to germinate at different temperatures are due to genetics, genetics x environment (GxE), or both. To assess the impact of GxE effect for germination time and rate, i.e., that the ranking of populations was conserved at the two temperatures tested, the low temperature phenotype was connected to high temperature phenotype for both species (Fig. 2; model 4). For GT, the relationship was significant for both species (Z-test, |z| > 3.95, p < 0.001). The adjusted R^2^ were of 0.86 and 0.51 for *B. rapa* and *B. oleracea* respectively. For GR, a significant link was observed for all accessions (|z| > 2.67, p < 0.007), except for wild accessions of *B. oleracea* (z = 0.97, p = 0.332). The adjusted R^2^ were of 0.76 and 0.08 for *B. rapa* and *B. oleracea* respectively.

**Figure 2.**
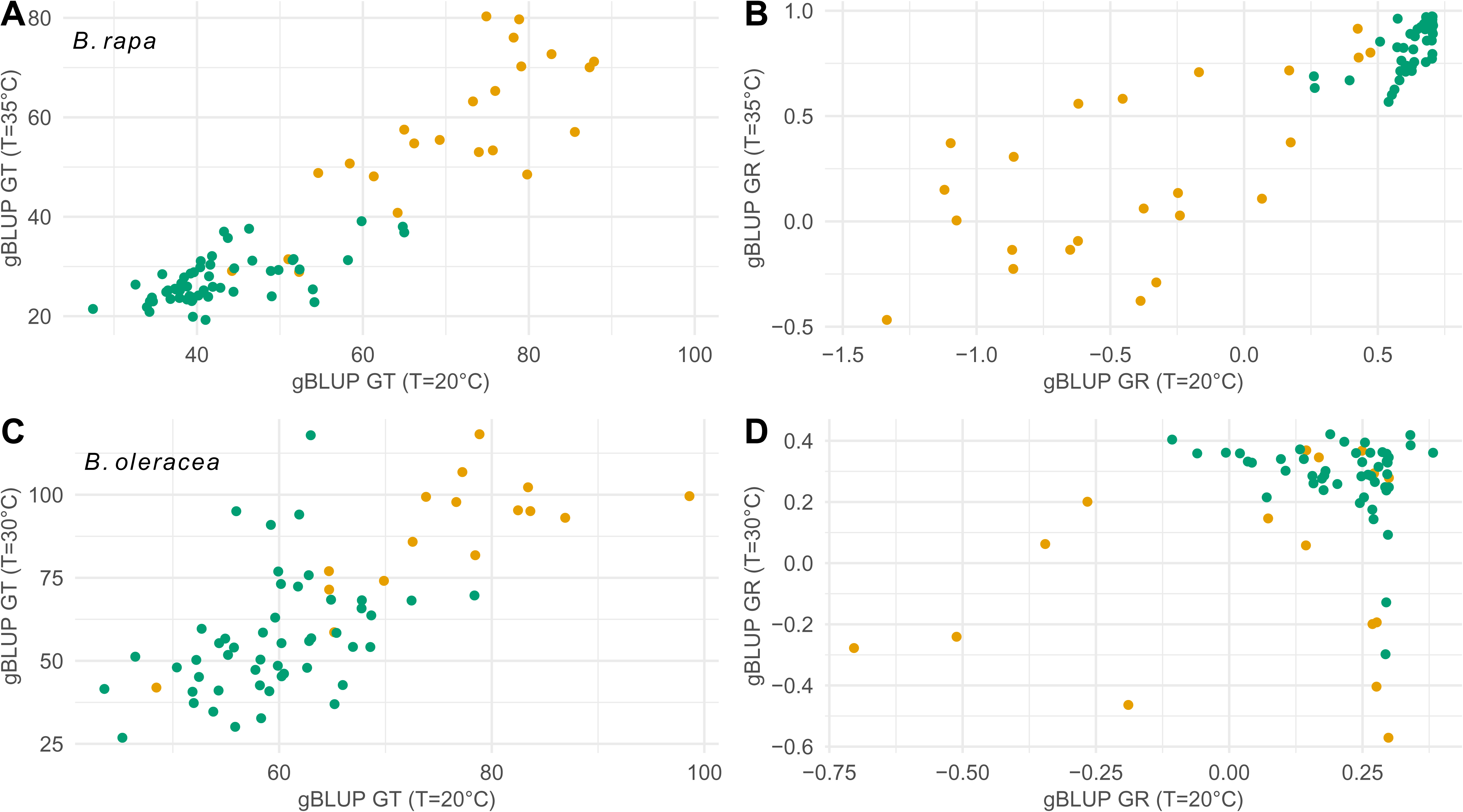
gBLUP of germination traits at 20°C as a function of the gBLUP of germination traits at 30/35°C. Wild accessions in orange; landraces in green. **(A)** BLUP of germination time of *B. rapa*. **(B)** BLUP of germination rate of *B. rapa*. **(C)** BLUP of germination time of *B. oleracea*. **(D)** BLUP of germination rate of *B. oleracea*.

### 4. The geographic provenance explained heat tolerance only for *B. oleracea* accessions

The GxE interactions in wild *B. oleracea* accessions could be explained by the environmental conditions at the sampling site. In fact, the latitude of the site, which was closely correlated with the mean annual temperature, only impacted the heat tolerance of *B. oleracea* accessions (Fig. 3). We tested this hypothesis in a manner analogous to Finlay-Wilkinson regression (Finlay & Wilkinson, 1963) with model (5). For *B. oleracea* accessions, southern accessions were more tolerant, as latitude had a significant effect on heat tolerance (Z-test, z < −2.56, p < 0.01) with an adjusted R ^2^ of 0.51. The effect of latitude was particularly pronounced among wild accessions (the effect of the latitude estimated at −0.017 for wild accessions, and −0.011 for landraces), where Spanish populations exhibited complete tolerance compared to French accessions. In contrast, no significant effect was observed for *B. rapa* wild accessions (z = −1.86, p = 0.064, with the effect of the latitude estimated at −0.006), nor for *B. rapa* landraces (z = −1.25, p = 0.215, with the effect of the latitude estimated at −0.009), albeit with an adjusted R^2^ of 0.62.

**Figure 3.**
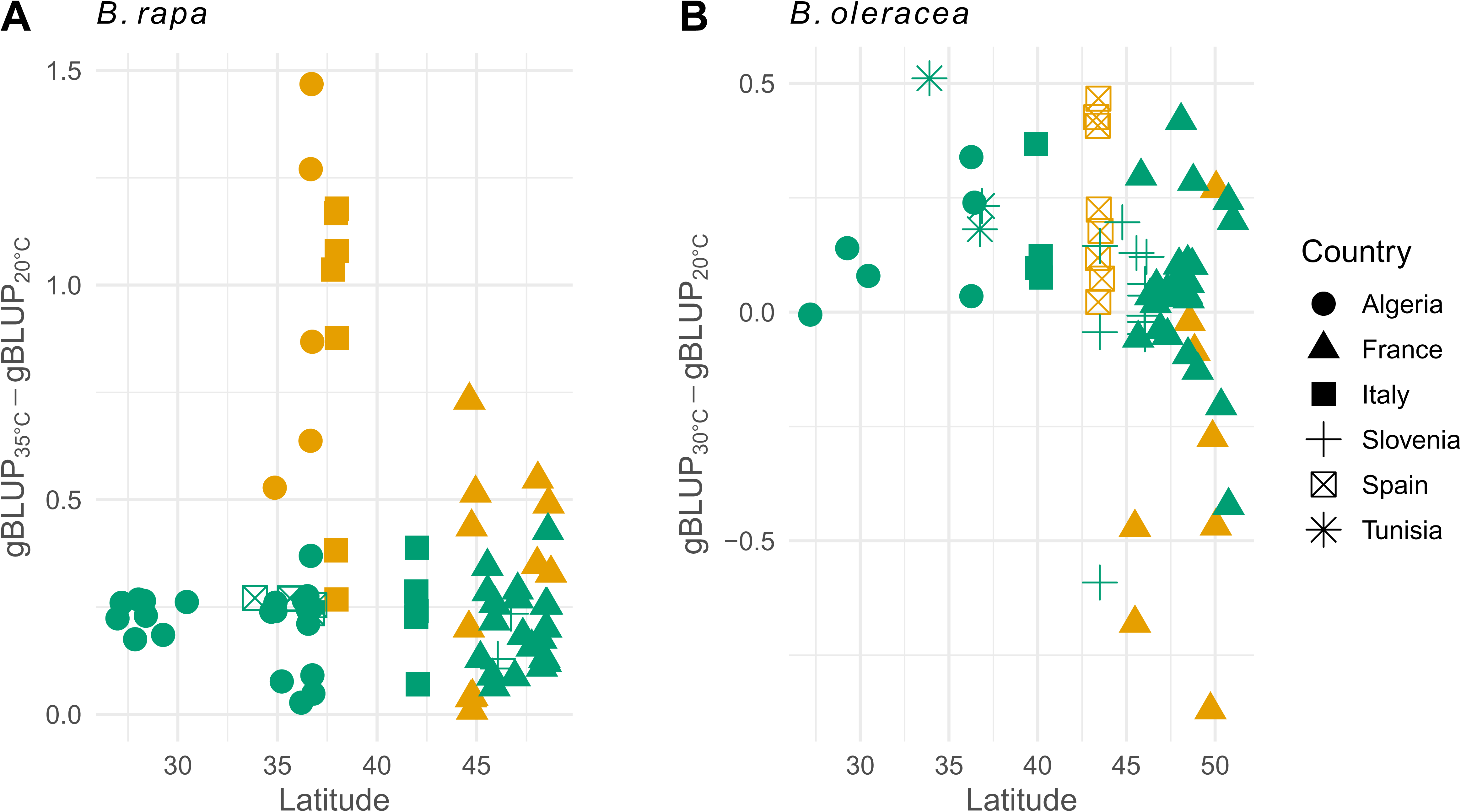
Heat tolerance (defined as the difference between the gBLUP at 30/35°C and that at 20°C) as a function of latitude. Wild accessions in orange; landraces in green. Provenance: Algeria: full circle; France: full triangle; Italy: full square; Slovenia: cross; Spain: crossed square; Tunisia: star. **(A)** Heat tolerance of *B. rapa*. **(B)** Heat tolerance of *B. oleracea*.

### 5. Germination was correlated with emergence only in *B. rapa* wild accessions

Germination is an essential component of seed vigor; however, in the context of agriculture, the ability to emerge is of greater importance. The genetic value of germination rate of *B. rapa* accessions measured in controlled conditions proved to be a reliable predictor of the emergence rate in the field for *B. rapa* accessions (Fig. 4; model 6), whereas it was not the case for *B. oleracea* accessions. For *B. rapa*, both the wild accessions and landraces showed a significant positive relationship (Z-test, z > 8.09, p < 0.001), with an adjusted R^2^ of 0.82. For *B. oleracea*, the relationship was not significant (|z| < 1.82, p > 0.078), with an adjusted R^2^ of 0.03.

**Figure 4.**
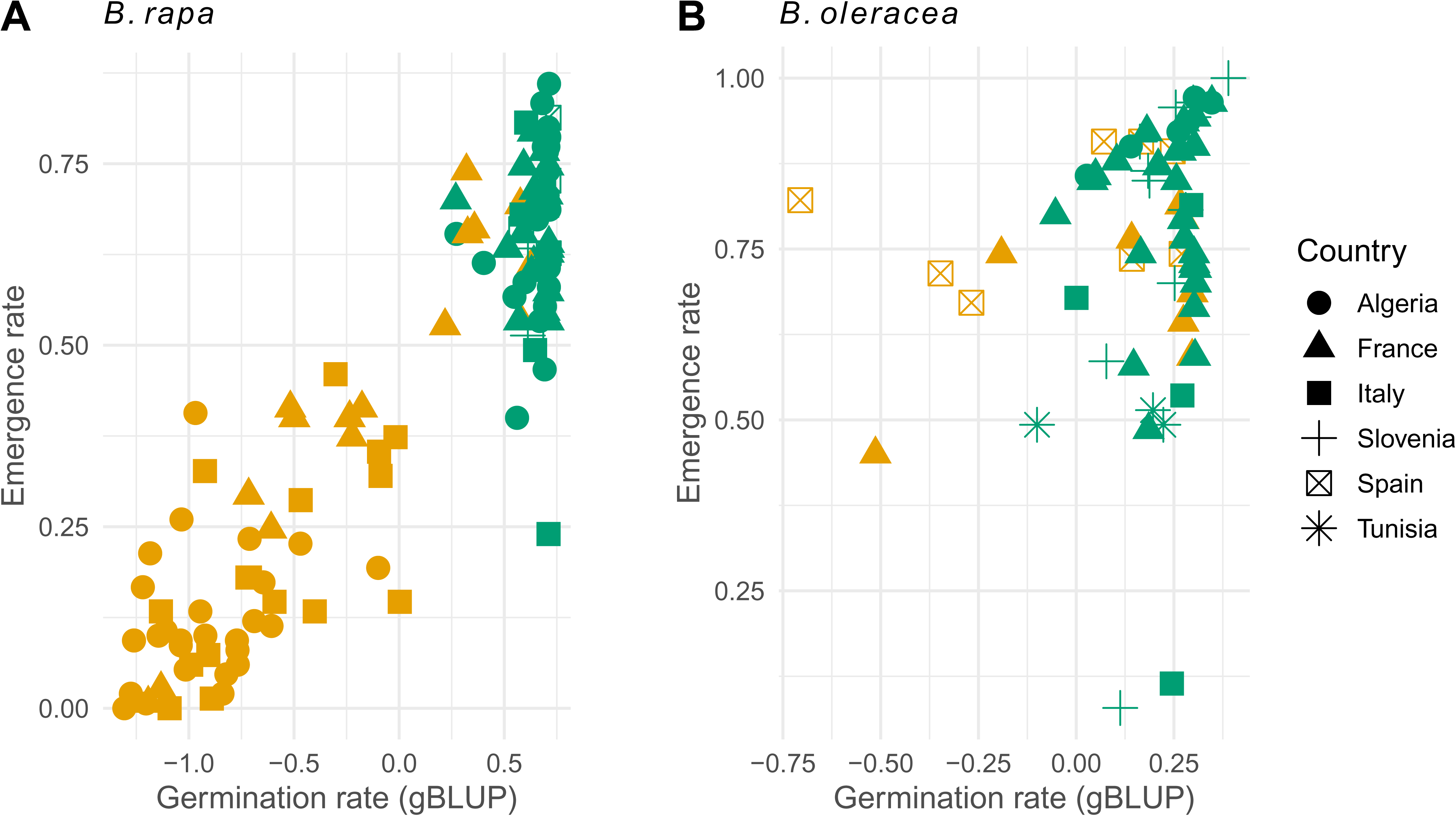
Emergence rate in field as a function of the gBLUP of germination rate in controlled conditions at 20°C. Wild accessions in orange; landraces in green. **(A)** Emergence rate of *B. rapa*. **(B)** Emergence rate of *B. oleracea*.

## Discussion

The extensive sampling of Mediterranean landscapes revealed a strong pattern of differentiation between the wild populations and the landraces, manifesting in both genetics and germination patterns. The elevated temperatures had a inhibitory or stimulatory effect on *B. oleracea* accessions depending on the provenance, whereas the *B. rapa* accessions generally benefited from the temperature increase. Comparing the gBLUP at both temperatures, results showed that an accession with superior germination capacity at 20°C also exhibited a robust capacity at warmer temperatures (30/35°C), indicating minimal GxE. Nevertheless, while all *B. rapa* accessions were heat tolerant regardless of the origin, the geographic provenance explained heat tolerance for *B. oleracea* accessions, wherein southern accessions demonstrated superior resilience to heat stress. Finally, the germination rate measured in laboratory settings proved to be a reliable predictor of emergence in the field only for *B. rapa*, suggesting that other factors are involved for *B. oleracea* emergence.

### 1. Different stories explains the different patterns of germination

The critical factor influencing germination appears to be the ‘type’ of the accessions, i.e., whether they were sampled in the wild or a cultivated landrace, regardless of the genetic clustering (e.g., French wild accessions of *B. rapa* cluster with landraces). It is important to note that a maternal effect is excluded as an explanatory factor, since all populations were multiplied for one generation in a common environment. As an illustration, germination processes of French *B. rapa* ‘wild’ populations are closest to that of other wild populations, with lower germination rates (GR) and higher germination times (GT), while being genetically closer to landraces. The literature suggests that these French populations may be feral (McAlvay et al., 2021). If this is the case, our results show that they tend to revert back to patterns akin to wild populations for germination, which is common, especially for dormancy (Ayal & Levy, 2005; Zhou et al., 2021). Similarly, the literature also suggests that all wild accessions of *B. oleracea* are more likely to be feral (Maggioni et al., 2020, 2024; Mittell et al., 2020; Cai et al., 2022; Saban et al., 2023), yet these populations displayed differences in GR and GT between ‘types’ of populations, albeit with weaker effects. This phenomenon suggests that, regardless of the ancestry of the populations, since there is less selection pressure for germination rate and time in the wild, bet-hedging strategies (Bewley, 1997) based on variable seed dormancy capacities might prevail. This adaptation is evident in *B. rapa* French populations, where dormancy – a trait typical of wild plants – was observed with high frequency (data not shown). In landraces, we can assume that such bet-hedging strategies have been selected against to favor homogeneity in the fields. This is consistent with the remarkably low variance observed for GR in landraces, which marks both the domestication bottleneck and the exhaustion of genetic variance by selection.

The different level of domestication between *B. rapa* and *B. oleracea* may also explain the lower differences between ‘types’ of *B. oleracea*: feralization in *B. oleracea* does not reduce germination rate as much as in *B. rapa* wild populations (i.e., French wild populations), possibly due to domestication fixation of high germination rate. This hypothesis is further supported by preliminary investigations that revealed no evidence of dormancy in *B. oleracea* accessions (data not shown), which is considered one of the first signs of domestication (Fuller & Allaby, 2009). Additionally, germination traits of cultivated lines of *B. napus* have been reported to be under strong selection (Laurençon et al., 2024). Furthermore, seed vigor is more closely correlated with vegetative growth than with performance at full reproductive maturity (Tekrony & Egli, 1991), which should have caused a strong selection for seed vigor even in cultivated lines of *B. oleracea* that are less often exploited for their seeds.

Interestingly, our findings indicate that the germination response to temperature is correlated with sampling site latitude (i.e., annual mean temperature) only for *B. oleracea*. The range of temperatures tested is a stress mainly for *B. oleracea* and not for *B. rapa*, with different intensity depending on the accession, which is reflected in the different levels of heat tolerance among the accessions. We expected accessions at lower latitudes to display higher adaptation to heat stress, resulting in higher heat tolerance, as they have been exposed to this selective pressure; this hypothesis was confirmed for *B. oleracea*, but not for *B. rapa*. The lower nucleotide diversity of *B. oleracea* than *B. rapa* (Saban et al., 2023) could indicate that stronger selection has acted on the former, perhaps imputable to a more pronounced domestication process or lower population sizes. In this study, the adaptation of landraces and feral populations to the local environment may be a signature of selection accounting for the environmental profiling, i.e., envirotyping (Xu, 2016; Colouer et al., 2019). In contrast, the lack of such a correlation in *B. rapa*, though a small effect on wild populations, suggests a strong selection of germination traits that are advantageous under a wide range of controlled agricultural environments, thereby reducing the influence of local climatic factors on germination (Purugganan & Fuller, 2009).

### 2. Genetic improvement for heat tolerance would be beneficial

Our findings show that there is genetic variance for germination traits, which suggests that those traits in these *Brassica* landraces could be selected and introgressed into commercial germplasms, which generally have an optimal germination temperature lower than 30-35°C (ISTA, 2024). The genotype x environment interaction was found to be minimal and there was significant genetic variances for GR and GT, which indicate that breeding can be conducted and worth promoting, given the unequal utilization of landraces across different countries (e.g., more common in Algeria than in France). For instance, *B. napus* seedlings show inhibited establishment at high temperatures (Zhang et al., 2015), indicating a potential benefit from integrating heat-tolerant traits from *B. rapa*. The tolerance of *B. rapa* landraces suggests that they will not suffer from future temperature rises and could be useful to improve commercial germplasms (Cochrane et al., 2015; Castillo-Lorenzo et al., 2019). Additionally, seed priming techniques could be explored to further enhance the germination performance of *B. rapa* under varying environmental conditions (Paparella et al., 2015). *B. oleracea* landraces experience heat as a stressor, significantly affecting their germination speed, although southern accessions show higher heat tolerance and could also prove useful for introgression into elite *B. oleracea* germplasms. Optimizing their use in breeding program is an active field of research; nowadays, pre-breeding and bridging populations act as a buffer to limit the linkage drag from wild materials and to maintain the level of performance of elite crop lines (Cowling et al., 2009; Sanchez et al., 2023).

The introgression of these traits is crucial for *B. napus* that could benefit from the germinative vigor of *B. rapa* landraces, especially if it also enhances emergence rates and crop field performance (Smith et al., 2000; Ghassemi-Golezani et al., 2010). Although our genome-wide association studies (GWAS) did not detect any major genes – consistent with the polygenic nature of germination traits (Hatzig et al., 2015; Bettey et al., 2000) – there is potential for improvement through phenomic and genomic selection (Laurençon et al., 2024). Nonetheless, the study has identified valuable sources of genetic diversity within *Brassica* populations, thereby highlighting the potential of expanding sampling efforts within these populations to circumvent the effect of population structure and increase the power of GWAS analyses (Tibbs Cortes et al., 2021). This would enable the identification of QTL underlying adaptation (Basnet et al., 2015), particularly those conferring resilience to temperature variation and improve the understanding of the genetic basis of adaptive traits (Huang & Han, 2014).

## Conclusion

The germination time and rate of an unexplored diversity of 25,280 seeds of *Brassica rapa* and 13,300 seeds of *Brassica oleracea* were compared at two temperature conditions. The results demonstrate that temperature elevation does not constitute a stressor for *B. rapa* accessions, which maintained high germination rates and short germination times at high temperatures, especially landraces. In contrast, some accessions of *B. oleracea* exhibited clear signs of stress when exposed to elevated temperatures, which in turn leads to a notable decline in their germination performance. Their level of heat tolerance depended on the geographical provenance, and in particular southern populations exhibited a higher degree of tolerance, likely due to their adaptation to higher annual mean temperatures. The integration of these heat-tolerant traits into commercial germplasms would enhance the resilience of *Brassica* species, ensuring their productivity and sustainability.

## Acknowledgement

We thank the Genetic Resource Centers BrACySol (https://www6.rennes.inrae.fr/igepp_eng/About-IGEPP/Platforms/BrACySol), especially V. Richer, S. Théréné, and S. Doré; and the Agricultural Institute of Slovenia (https://www.kis.si/en/) for providing seeds from different landraces. We thank the GenOuest bioinformatics platform (https://www.genouest.org/), as well as the Plant imaging platform PHENOTIC in Angers (https://phenotic.hub.inrae.fr/). We also extend our appreciation to the staff who provided care for the plant material, particularly L. Charlon, J-P. Constantin, and F. Letertre; P. Glory and P. Vallée for field experiments; and to M. Gautier and M. Delmond for the SNP calling.

## Contributions

MHW, CF, DS, and AMC designed the study; FG and KL generated Illumina data; MHW, AD performed experiments and contributed data; MT, MHW, EC, LG, and LM analyzed the data; MT, LG, and MHW drafted the manuscript. All authors read and approved the final version of the manuscript.

## Data availability

The short-read resequencing data that support the findings are available in NCBI’s sequence read archive (SRA) under the project PRJNA1174687. The genetic files (bed, bim, fam) and the raw phenotype dataset is freely available at: “Tiret, Mathieu, 2025, Replication Data for: An unexplored diversity for adaptation of germination to high temperatures in Brassica species, https://doi.org/10.57745/RIOMJ8”. Seeds of the accessions are available at the following Genetic Resource Centers: BrACySol for French accessions, de Los Campos seed genebank (ESP003) for Spanish accessions, and Agricultural Institute of Slovenia for Slovenian accessions. The unique accession numbers are detailed in Table S3. For inquiries regarding Algerian, Italian, and Tunisian accessions, please refer to the contact specified in Table S3.

