## Supplementary material for "An unexplored diversity for adaptation of germination to high temperatures in *Brassica* species"

### Supplementary materials

|  | T° | B. rapa | | B. oleracea | |
| --- | --- | --- | --- | --- | --- |
|  |  | Wild | Landrace | Wild | Landrace |
| GR | 20°C | 0.22 | 0.10 | 0.19 | 0.08 |
|  | 30/35°C | 0.19 | 0.13 | 0.13 | 0.08 |
| GT | 20°C | 0.13 | 0.18 | 0.33 | 0.13 |
|  | 30/35°C | 0.19 | 0.11 | 0.29 | 0.34 |

**Table S1**. Heritability of germination time (GT) and rate (GR) estimated in the models (1c) and (3c).

|  | T° | B. rapa | | B. oleracea | |
| --- | --- | --- | --- | --- | --- |
|  |  | Wild | Landrace | Wild | Landrace |
| GR | 20°C | 0.29 | 0.02 | 0.16 | 0.02 |
|  | 30/35°C | 0.25 | 0.03 | 0.13 | 0.02 |
| GT | 20°C | 212.12 | 104.99 | 197.17 | 74.88 |
|  | 30/35°C | 309.05 | 38.44 | 514.25 | 467.69 |

**Table S2**. Additive variances of germination time and rate estimated in the models (1c) and (3c).

**Table S3**. List of the accessions, their accession numbers, country of origin, type (Landrace or Wild), and reference contact.

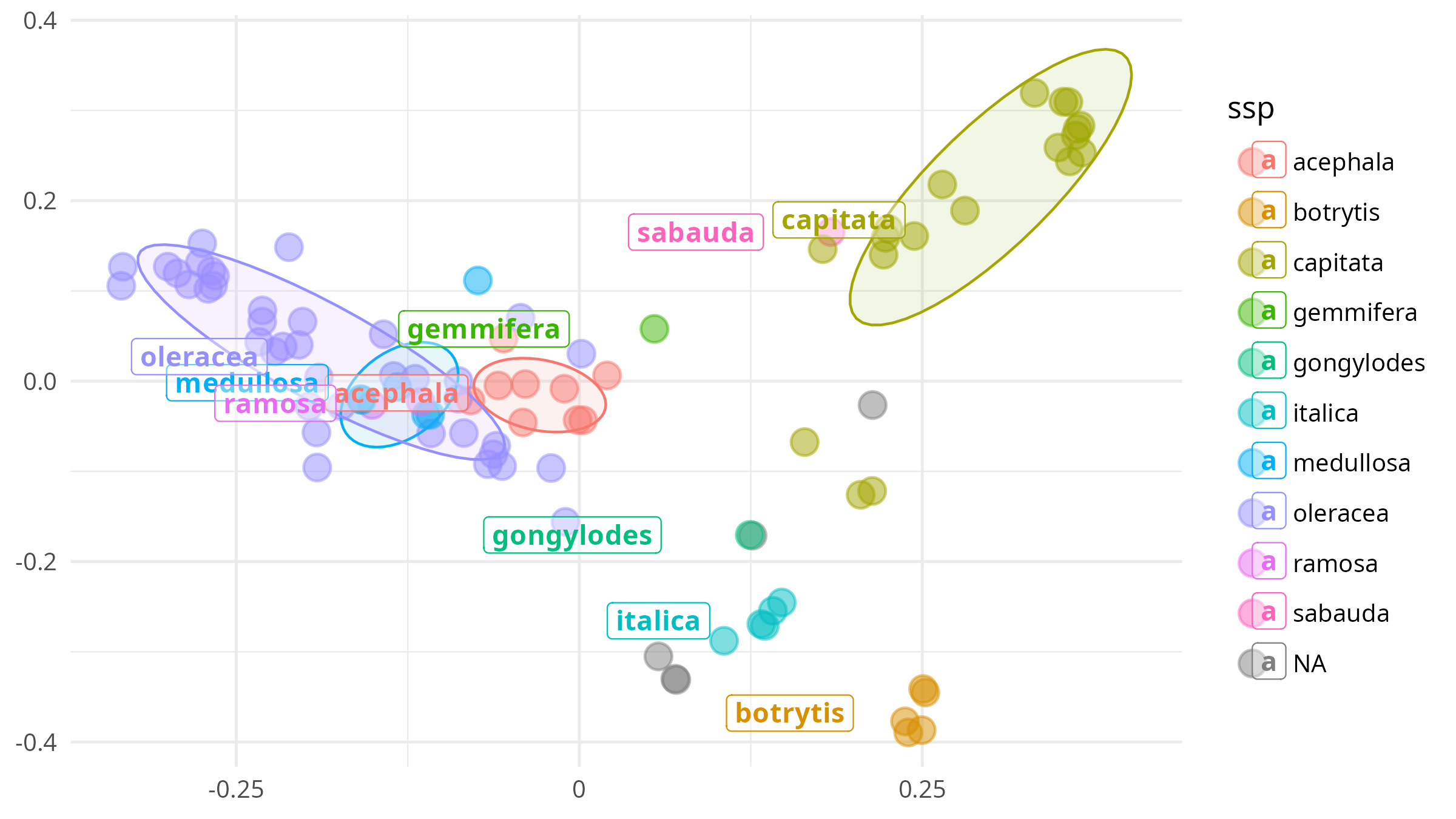
**Figure S1**. MDS on the genotypes of *B. oleracea* accessions, colored by morphotypes. The morphotype of four landraces from the south of Algeria was unknown (denoted NA).

#### A. GWAS

Using the gBLUP of the models (1c) and (3c), we estimated the effect sizes of SNPs in a GWAS with GCTA v1.94.1 with default parameters. The confidence on these GWAS is yet too low because of the sample size (less than 200 genotypes).

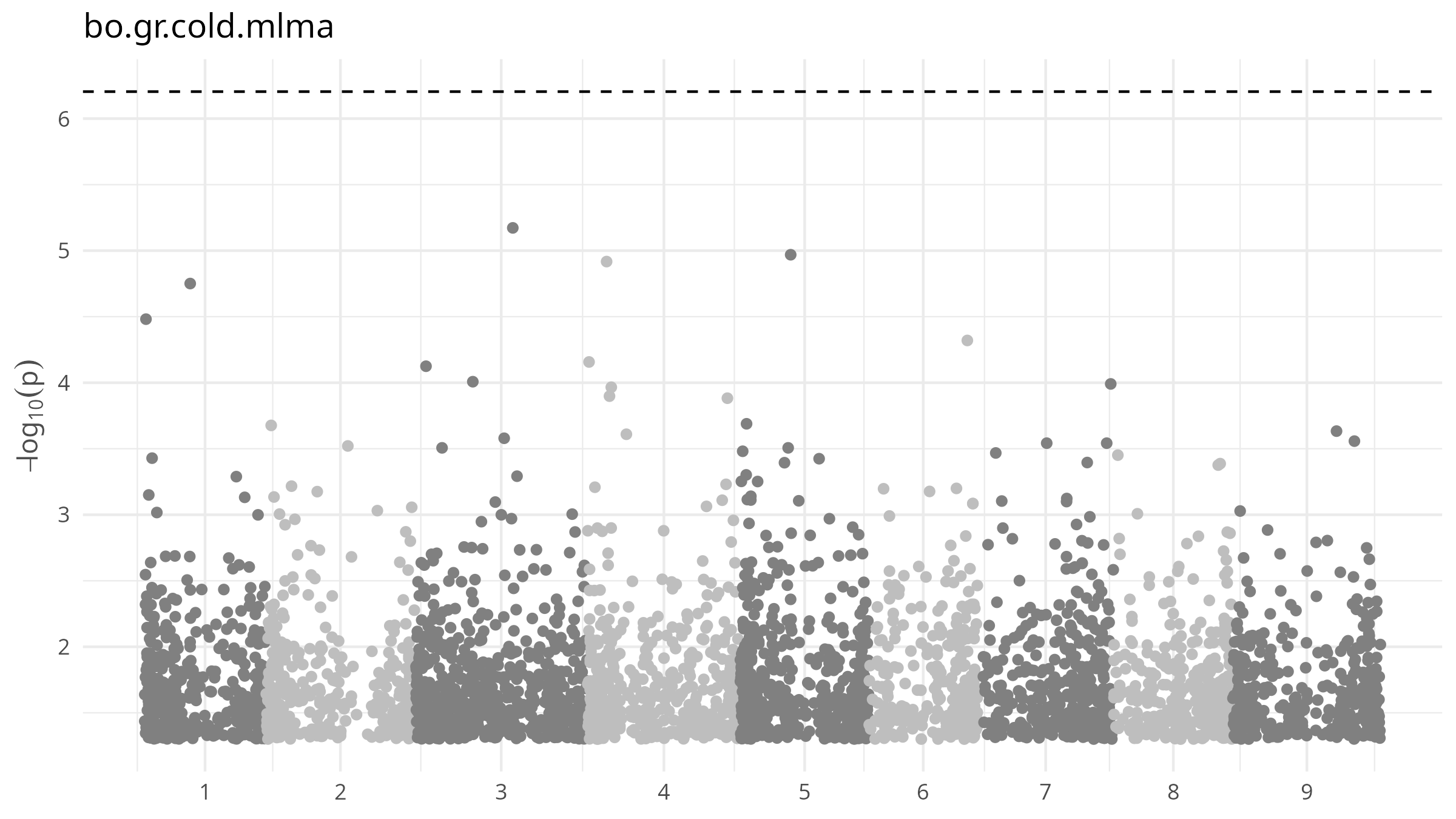
**Figure S2**. GWAS on the gBLUP of GR at 20°C on *B. oleracea* accessions. The dotted horizontal line is the Bonferroni correction threshold.

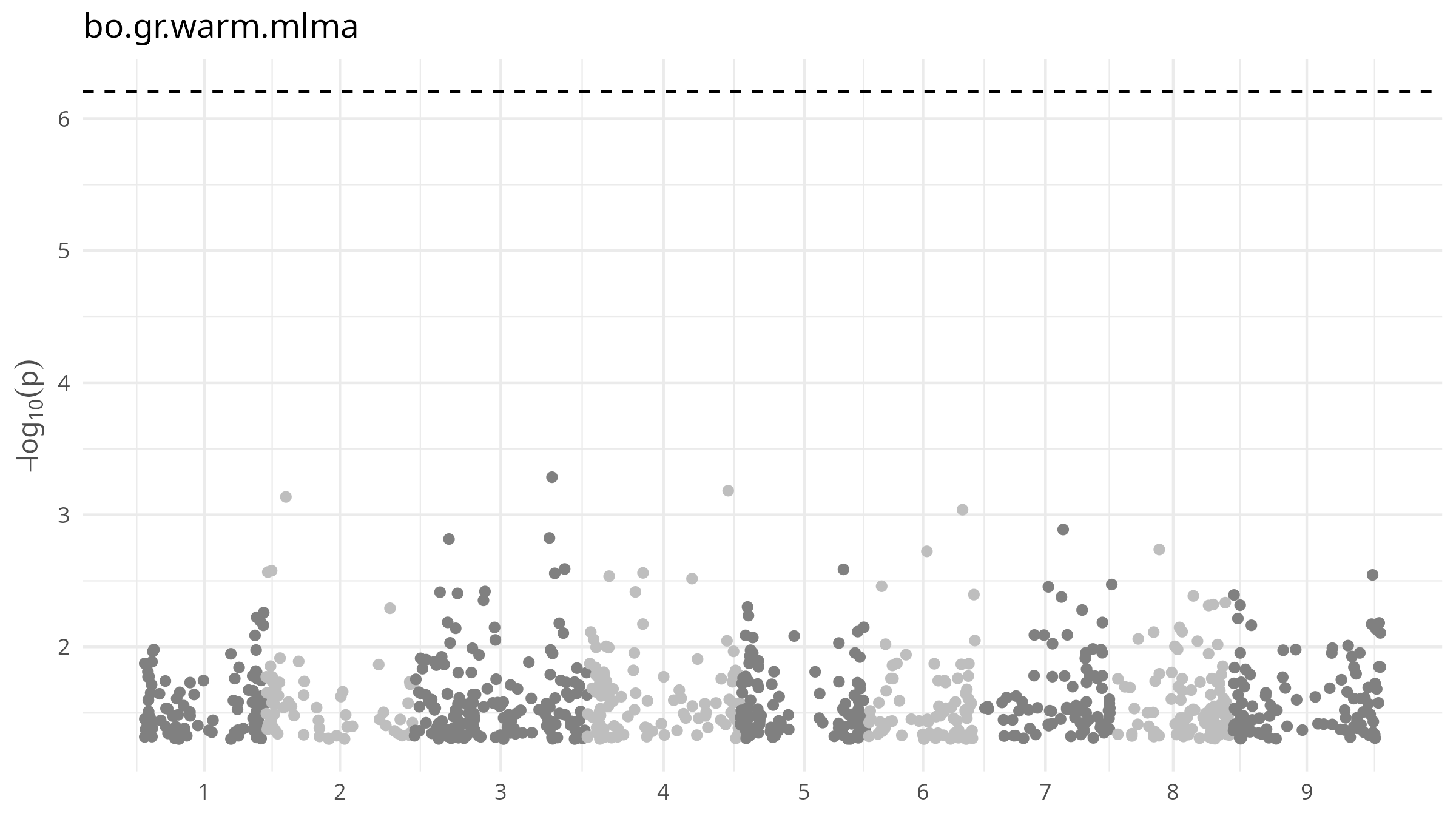
**Figure S3**. GWAS on the gBLUP of GR at 30°C on *B. oleracea* accessions.

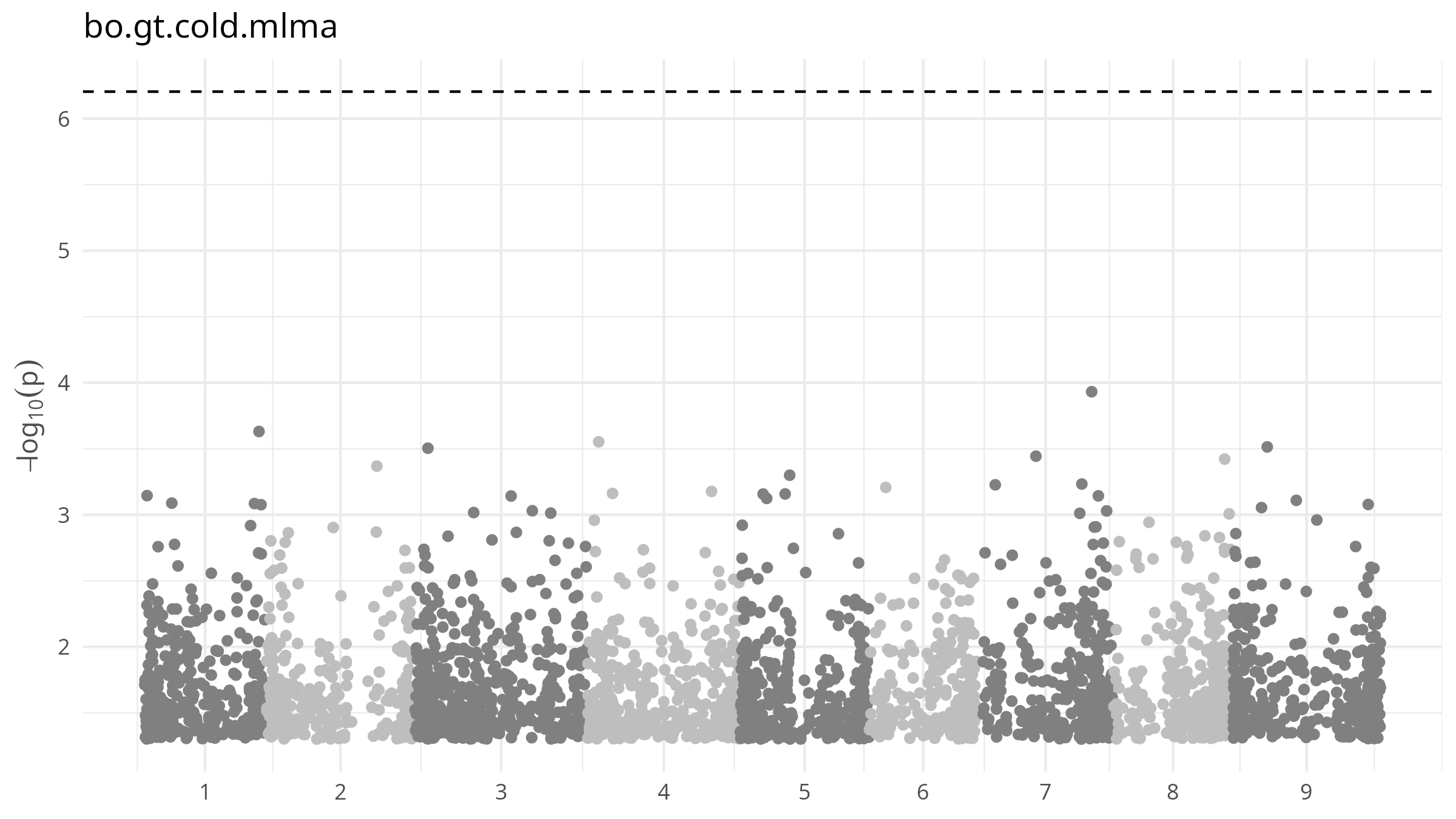
**Figure S4**. GWAS on the gBLUP of GT at 20°C on *B. oleracea* accessions.

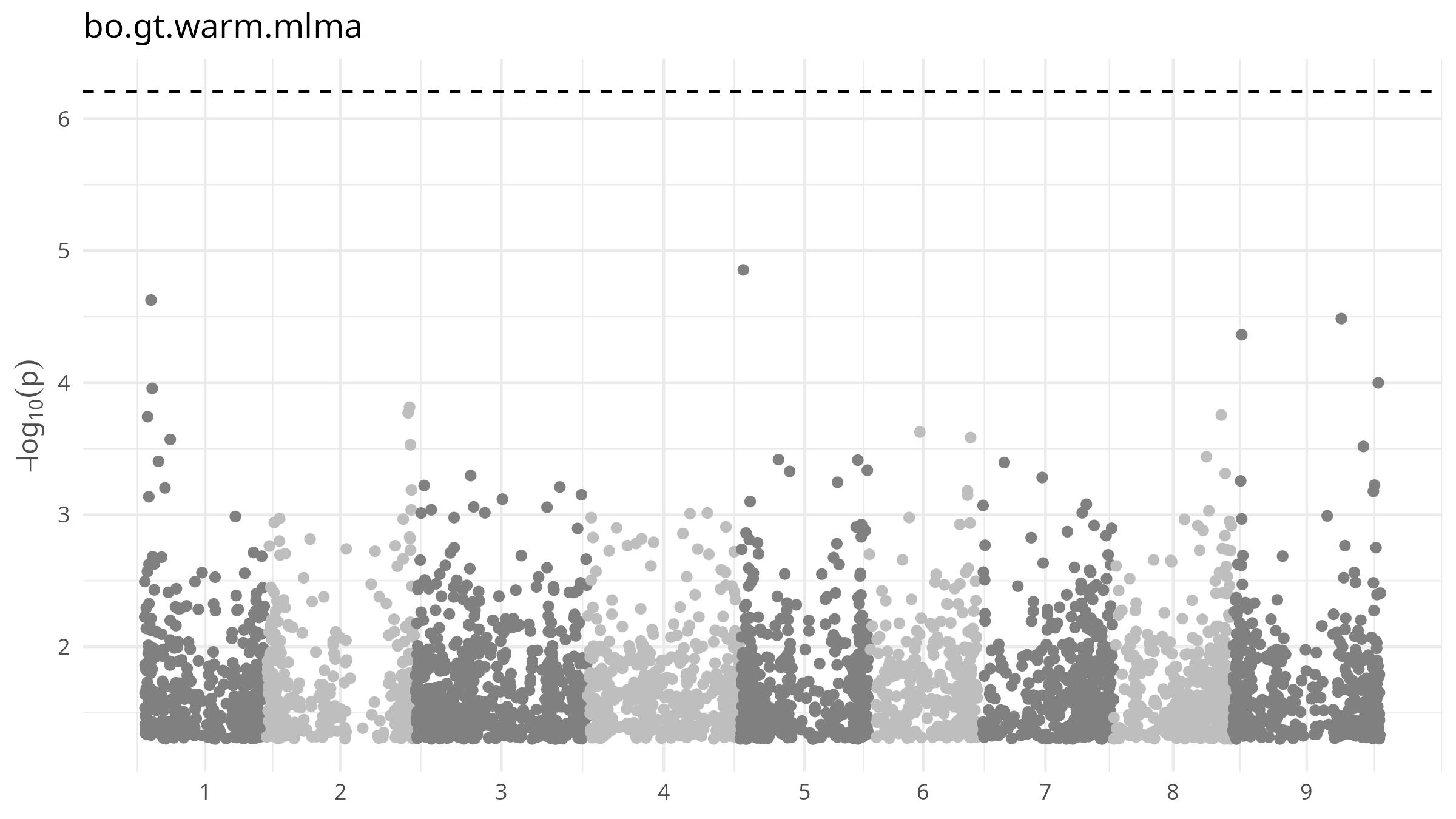
**Figure S5**. GWAS on the gBLUP of GT at 30°C on *B. oleracea* accessions.

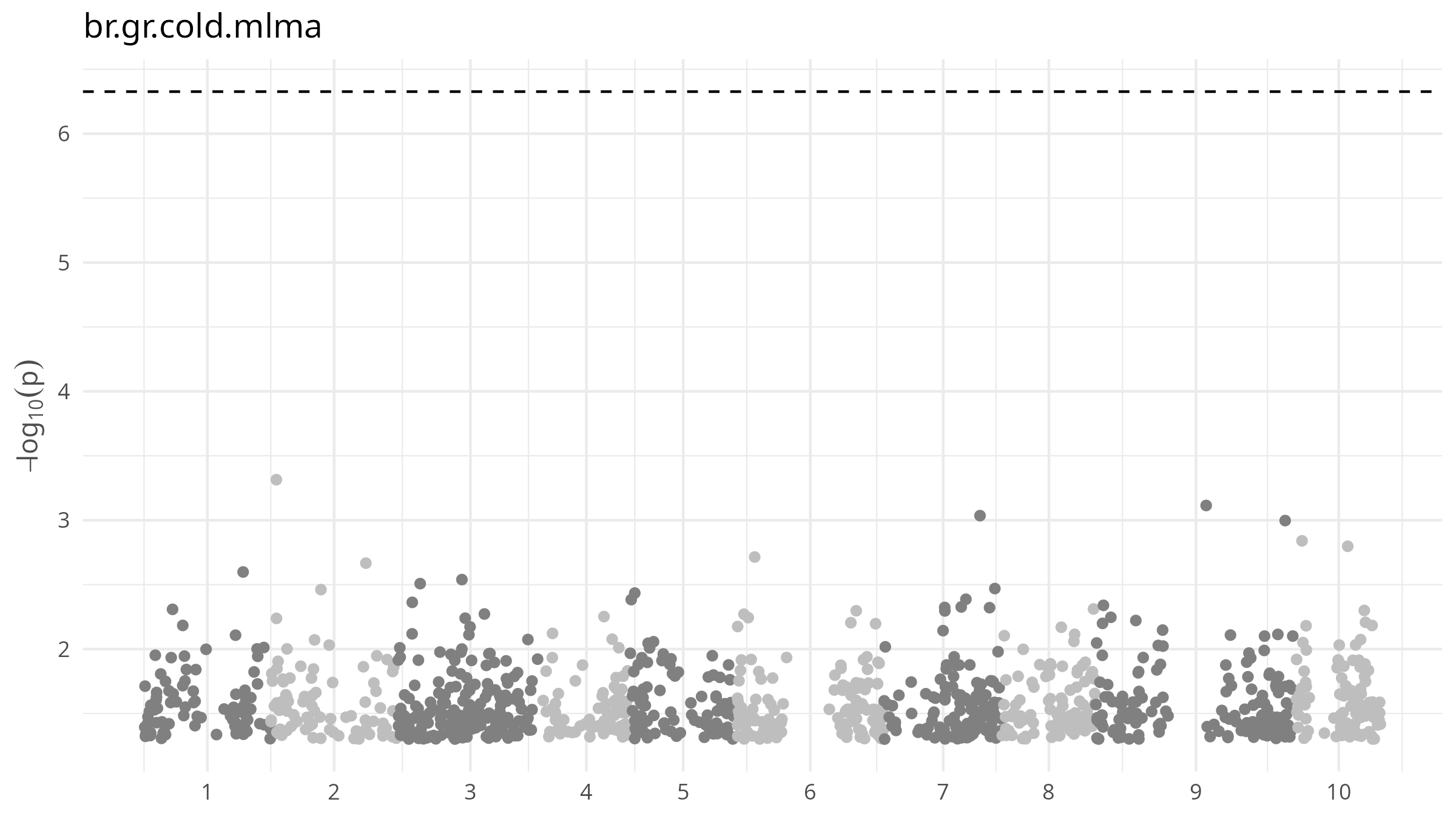
**Figure S6**. GWAS on the gBLUP of GR at 20°C on *B. rapa* accessions.

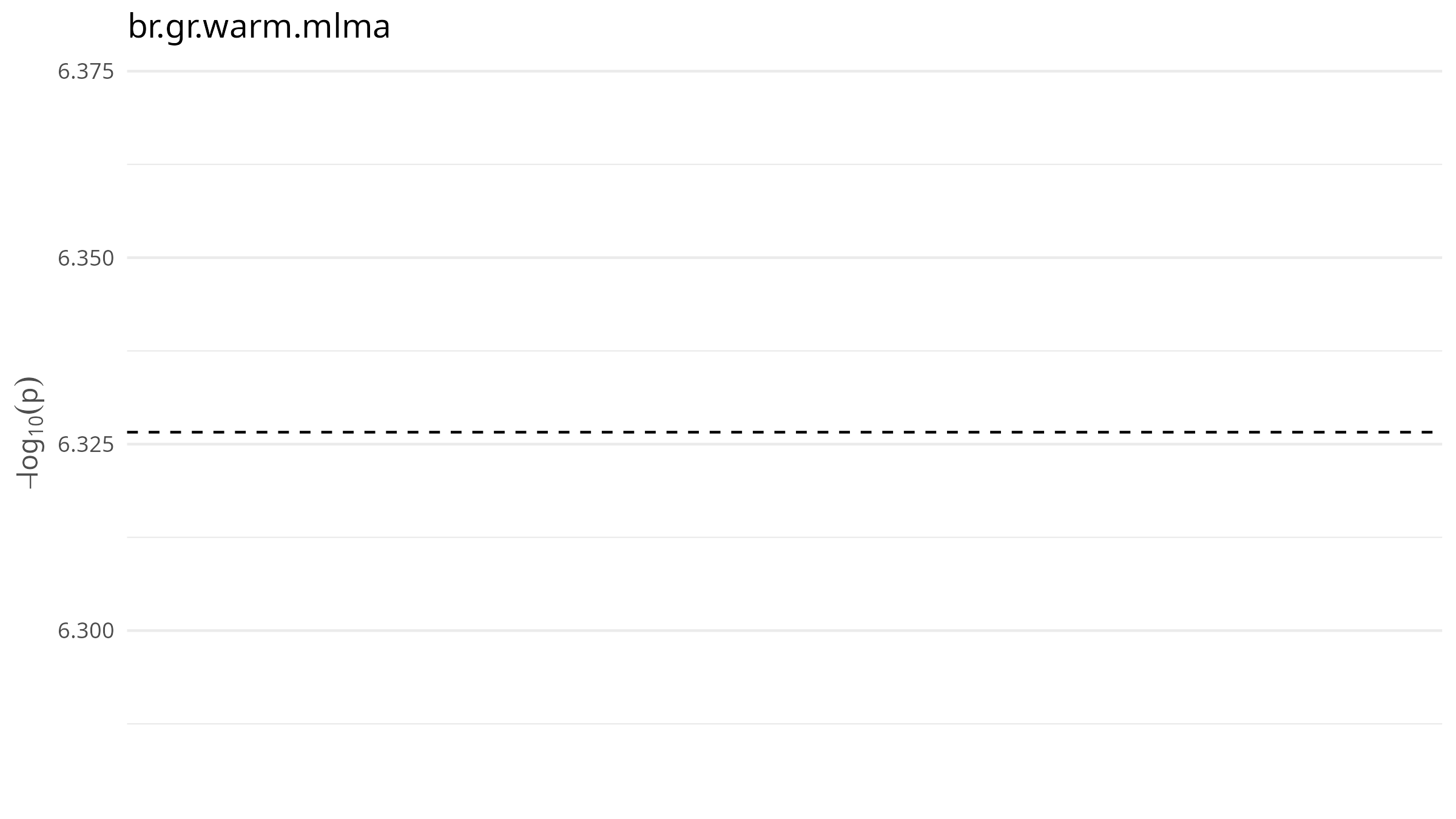
**Figure S7**. GWAS on the gBLUP of GR at 35°C on *B. rapa* accessions. The software (GCTA) did not converge.

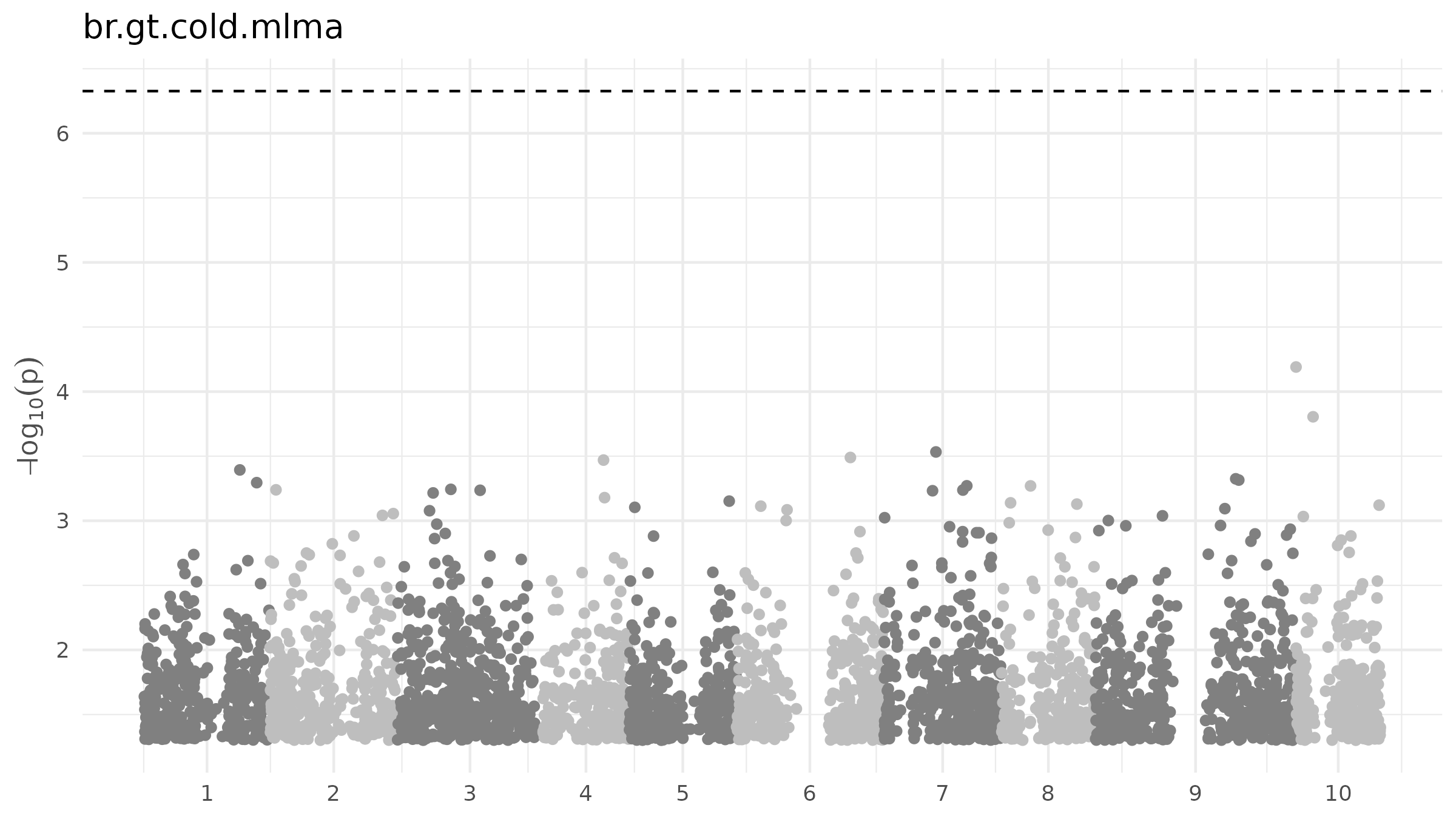
**Figure S8**. GWAS on the gBLUP of GT at 20°C on *B. rapa* accessions.

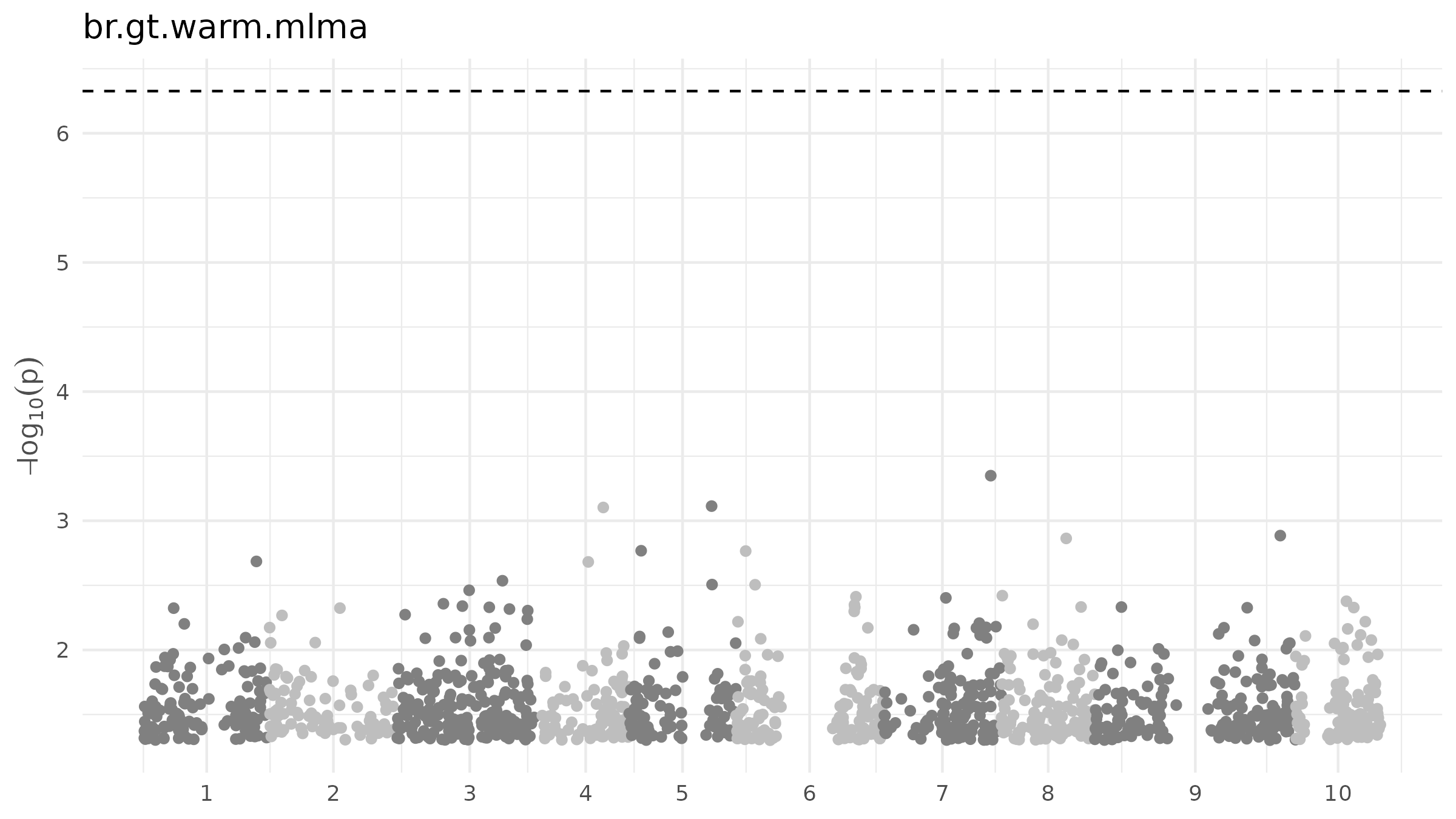
**Figure S9**. GWAS on the gBLUP of GT at 35°C on *B. rapa* accessions.
